# CD14 Blockade Does Not Improve Outcomes of Deep Vein Thrombosis Following Inferior Vena Cava Stenosis in Mice

**DOI:** 10.1101/2024.01.10.575099

**Authors:** Nilesh Pandey, Harpreet Kaur, Lakshmi Chandaluri, Sumit Kumar Anand, Himanshu Chokhawala, Tarek Magdy, Karen Y Stokes, A Wayne Orr, Oren Rom, Nirav Dhanesha

## Abstract

**Background:** Neutrophil-mediated persistent inflammation and neutrophil extracellular trap formation (NETosis) promote deep vein thrombosis (DVT). CD14, a co-receptor for toll-like receptor 4 (TLR4), is actively synthesized by neutrophils, and the CD14/TLR4 signaling pathway has been implicated in proinflammatory cytokine overproduction and several aspects of thromboinflammation. The role of CD14 in the pathogenesis of DVT remains unclear.

**Objective:** To determine whether CD14 blockade improves DVT outcomes.

**Methods:** Bulk RNA sequencing and proteomic analyses were performed using isolated neutrophils following inferior vena cava (IVC) stenosis in mice. DVT outcomes (IVC thrombus weight and length, thrombosis incidence, neutrophil recruitment, and NETosis) were evaluated following IVC stenosis in mice treated with a specific anti-CD14 antibody, 4C1, or control antibody.

**Results:** Mice with IVC stenosis exhibited increased plasma levels of granulocyte colony-stimulating factor (G-CSF) along with a higher neutrophil-to-lymphocyte ratio and increased plasma levels of cell-free DNA, elastase, and myeloperoxidase. Quantitative measurement of total neutrophil mRNA and protein expression revealed distinct profiles in mice with IVC stenosis compared to mice with sham surgery. Neutrophils of mice with IVC stenosis exhibited increased inflammatory transcriptional and proteomic responses, along with increased expression of CD14. Treatment with a specific anti-CD14 antibody, 4C1, did not result in any significant changes in the IVC thrombus weight, thrombosis incidence, or neutrophil recruitment to the thrombus.

**Conclusion:** The results of the current study are important for understanding the role of CD14 in the regulation of DVT and suggest that CD14 lacks an essential role in the pathogenesis of DVT following IVC stenosis.

## 1. INTRODUCTION

Venous thromboembolism (VTE), which encompasses deep vein thrombosis (DVT) and pulmonary embolism (PE), affects an estimated 900,000 people in the United States. VTE poses significant morbidity and mortality risks, and imposes a substantial economic burden on individuals, healthcare systems, and society. Despite recent advances in therapeutic management, the incidence of VTE continues to rise because of increased life expectancy and the prevalence of conditions that predispose patients to VTE.^1-4^ A better understanding of the molecular mechanisms that facilitate venous thrombus propagation may, therefore, lead to the development of novel and safe therapeutics for VTE.

Patients with VTE exhibit an acute proinflammatory response, characterized by systemic leukocytosis and increased inflammatory cytokines.^5-7^ Elevated leukocytes, particularly neutrophils, is strongly associated with an increased risk of VTE and mortality.^8-10^ At the venous shear rate, neutrophils promote thrombus growth through several mechanisms, such as the release of neutrophil extracellular traps (NETs), secretion of inflammatory mediators, and promotion of endothelial cell activation.^1,7^ Neutrophils-mediated persistent inflammation and NETosis may simultaneously affect coagulation and fibrinolysis, resulting in venous thrombus propagation, and contributing to the development of subsequent complications. Despite the clear involvement of neutrophils in the pathogenesis of DVT, alterations in transcriptome and proteome following DVT have not yet been evaluated.

To gain insight into the regulation of neutrophils in response to DVT, we analyzed plasma levels of granulocyte colony-stimulating factor (G-CSF), blood neutrophil counts, and performed unbiased RNA sequencing (RNA-seq) and proteomics of primary mouse neutrophils following inferior vena cava (IVC) stenosis. IVC stenosis is the gold standard murine model that mimics the conditions of disturbed flow leading to DVT in humans. We identified distinct transcriptional and proteomic profiles in neutrophils from mice with IVC stenosis compared to sham-surgery mice. Strikingly, CD14 was aberrantly overexpressed in the bone marrow and peripheral blood neutrophils of mice with DVT. CD14, a co-receptor for toll-like receptor 4 (TLR4), is mainly expressed in monocytes and macrophages. CD14 is also actively synthesized by neutrophils, and the CD14/TLR4 signaling pathway has been implicated in proinflammatory cytokine overproduction and several aspects of thromboinflammation.^11,12^ Numerous pre-clinical studies and early phase clinical studies suggest that CD14-blockade with a specific monoclonal antibody may exert protective effects in diseases involving excessive activation of innate immunity by reducing sepsis-induced coagulopathy and inflammation.^11-15^ Here, we hypothesized that CD14 blockade could reduce DVT severity following IVC stenosis. Contrary to our hypothesis, anti-CD14 antibody treatment did not improve DVT outcomes. These findings are important for understanding the role of CD14 in the regulation of venous thrombus propagation and suggest that CD14 lacks an essential role in the pathogenesis of DVT following IVC stenosis.

## 2. METHODS

Additional methods are available in supplemental material.

### 2.1 Mice

All animal procedures were approved by the Institutional Animal Care & Use Committees of Louisiana State University Health Sciences Center-Shreveport (P-22-023). Ten to twelve old male C57Bl/6J (WT) mice were purchased from Jackson Lab.

### 2.2 Inferior vena cava (IVC) stenosis model

The mouse IVC stenosis model of DVT was performed in mice, as previously described.^4,16-18^ Only male mice were used for this model, as ligation in female mice may result in necrosis of the reproductive organs.^17,19^ Briefly, a midline laparotomy was made, IVC side branches were first ligated. For stenosis, a space holder (30-gauge) was positioned on the outside of the exposed IVC, and a permanent narrowing ligature was placed below the left renal vein. Next, the needle was removed to restrict blood flow to 80-90%. IVC thrombi were harvested 48-hour post-stenosis, detached from the vessel wall, dried, and weighed in a microbalance, and imaged. The operator was blinded with respect to the treatment of mice.

### 2.3 Statistical analysis

For analysis, GraphPad Prism software (9.4.1) was used. Normality and equal variance were tested using the Shapiro-Wilk and Bartlett’s test, respectively. Normally distributed data were analyzed by Student’s t-test or two-way ANOVA followed by Sidak’s multiple comparisons test, and non-normally distributed data were analyzed using the Mann Whitney test (for two-group) or non-parametric two-way ANOVA followed by Fischer’s LSD test. Thrombosis incidence data were analyzed using the Fisher’s exact test. The results were considered significant at P<0.05.

## 3. RESULTS AND DISCUSSION

### Mice with experimental DVT exhibit increased plasma G-CSF levels, neutrophilia, and neutrophil hyperactivation

G-CSF plays a fundamental role in granulopoiesis and in activation of mature neutrophils. Increased plasma G-CSF levels and higher blood neutrophil counts have been reported in several inflammatory conditions.^20-22^ To evaluate the effect of experimental DVT on plasma G-CSF levels, we subjected wild-type (WT) mice to IVC stenosis or sham surgery. We evaluated plasma G-CSF levels at 3, 6, and 24-hour post IVC stenosis and observed significantly increased plasma G-CSF levels at all time points tested in mice with IVC stenosis compared to mice with sham surgery **(Figure 1A-B)**. Analysis of complete blood count data revealed that mice with DVT exhibited significantly increased blood neutrophils, reduced blood lymphocytes **(Supplemental Figure 1A-B)**, and increased neutrophil-to-lymphocyte ratio **(Figure 1C)**, while platelet and monocyte counts did not significantly change **(Supplemental Figure 1C-D)**. These changes in blood neutrophil counts were also associated with increased plasma levels of cell-free DNA **(Figure 1D)** and increased plasma levels of neutrophil granule proteins such as elastase and myeloperoxidase (MPO) **(Figure 1 E-F)** in mice with DVT. These data suggest that mice with experimental DVT exhibit increased plasma G-CSF levels, neutrophilia, and neutrophil hyperactivation and are in agreement with previous reports suggesting increased G-CSF levels correlate with NET-associated thrombosis in animal models.^23,24^

**Figure 1:**
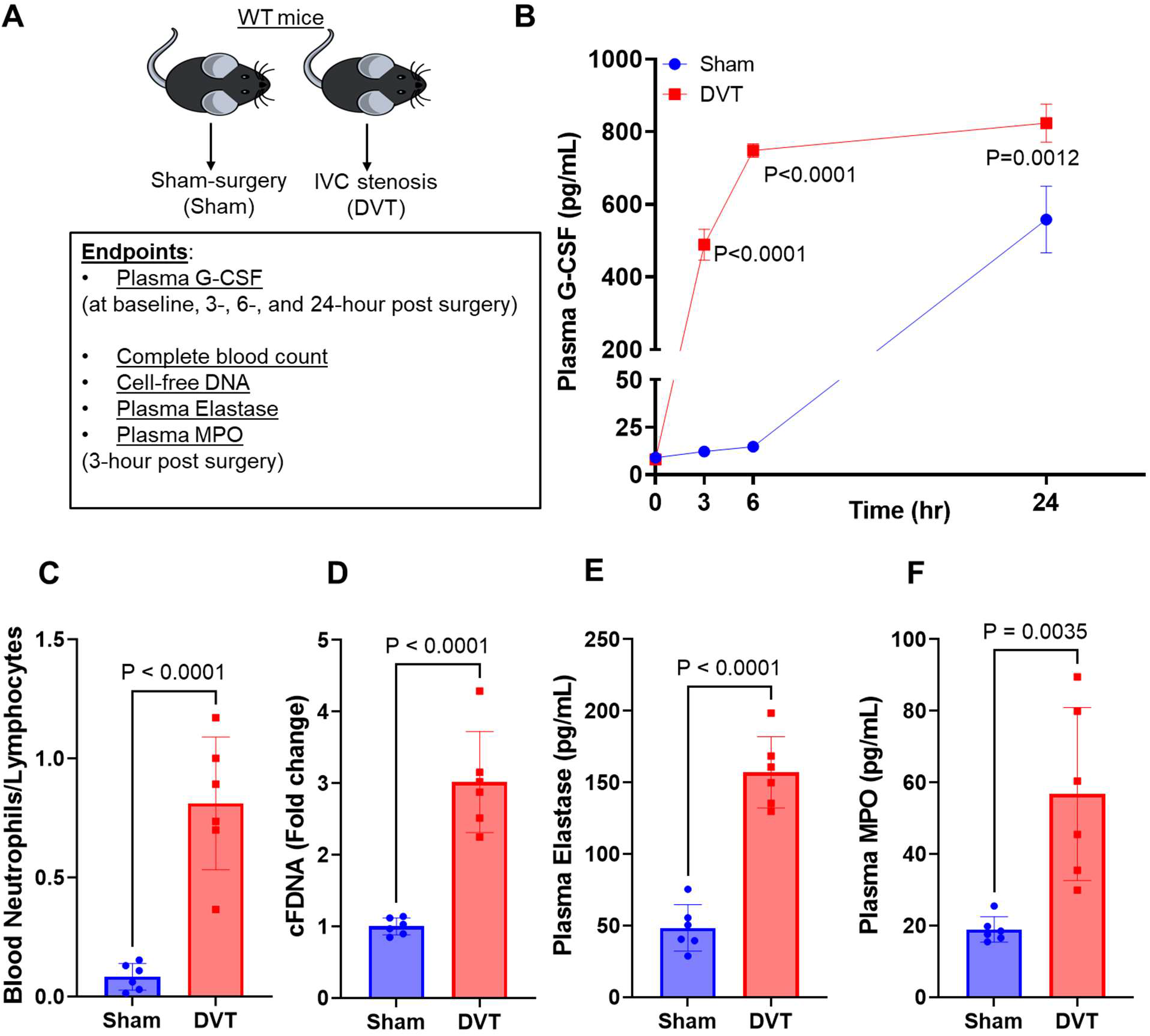
Mice with experimental DVT exhibit increased plasma G-CSF levels, neutrophilia, and neutrophil hyperactivation. (A) Schematic of experimental design. (B) Plasma G-CSF levels in mice with IVC stenosis and with sham-surgery at indicated time points. Blood was collected 3-hr post-surgery from mice with sham-surgery and mice with IVC stenosis for analysis of (C) Blood neutrophil/lymphocyte ratio, (D) cell-free DNA, (E) plasma elastase, and (F) plasma MPO. All data are from male mice and are mean ± SEM and analyzed by 2-way repeated measures ANOVA (Kruskal-Wallis test) followed by Fisher’s LSD test (B), or an unpaired Student t test (C-F); n = 3 (B) and n = 6 (C-F).

### Transcriptomics and proteomics analysis of post-DVT neutrophils revealed aberrant overexpression of CD14

To evaluate post-DVT transcriptional responses, we performed bulk RNA-seq of neutrophils isolated from mice with IVC stenosis and compared them to those from mice subjected to sham surgery **(Figure 2A)**. Principal component analysis (PCA) of neutrophil RNA-seq 3-hour post-surgery showed distinct transcriptional profiles in mice with IVC stenosis and mice with sham-surgery **(Figure 2B)**. Analysis of differential gene expression (P-adj < 0.01) revealed a total of 1454 differentially expressed genes with 372 genes upregulated, and 1082 genes downregulated in neutrophils from mice with IVC stenosis relative to mice with sham surgery **(Figure 2C)**. Hierarchical clustering of gene expression revealed gene clusters that were highly upregulated in mice with IVC stenosis **(Supplemental Figure 2)**. The top 20 significantly altered genes are shown in **Figure 2D**. Genes found to be upregulated in mice with IVC stenosis were proinflammatory (*Cd14, Nos2, Saa3, Acod1, Plscr1*), anti-apoptotic (*Muc1, Bcl3, Snai1,Bcl2, Ddit4, CD47*), and involved in chemotaxis (*Cxcl3, Cxcl2, Ccl2, Pded4d*) **(Figure 2E)**. Gene ontology (GO) enrichment revealed that highly upregulated genes in mice with IVC stenosis corresponded to biological processes involved in translation, ribosome biogenesis, response to cytokines, intrinsic apoptotic pathways, and inflammatory responses **(Figure 2F)**.

**Figure 2:**
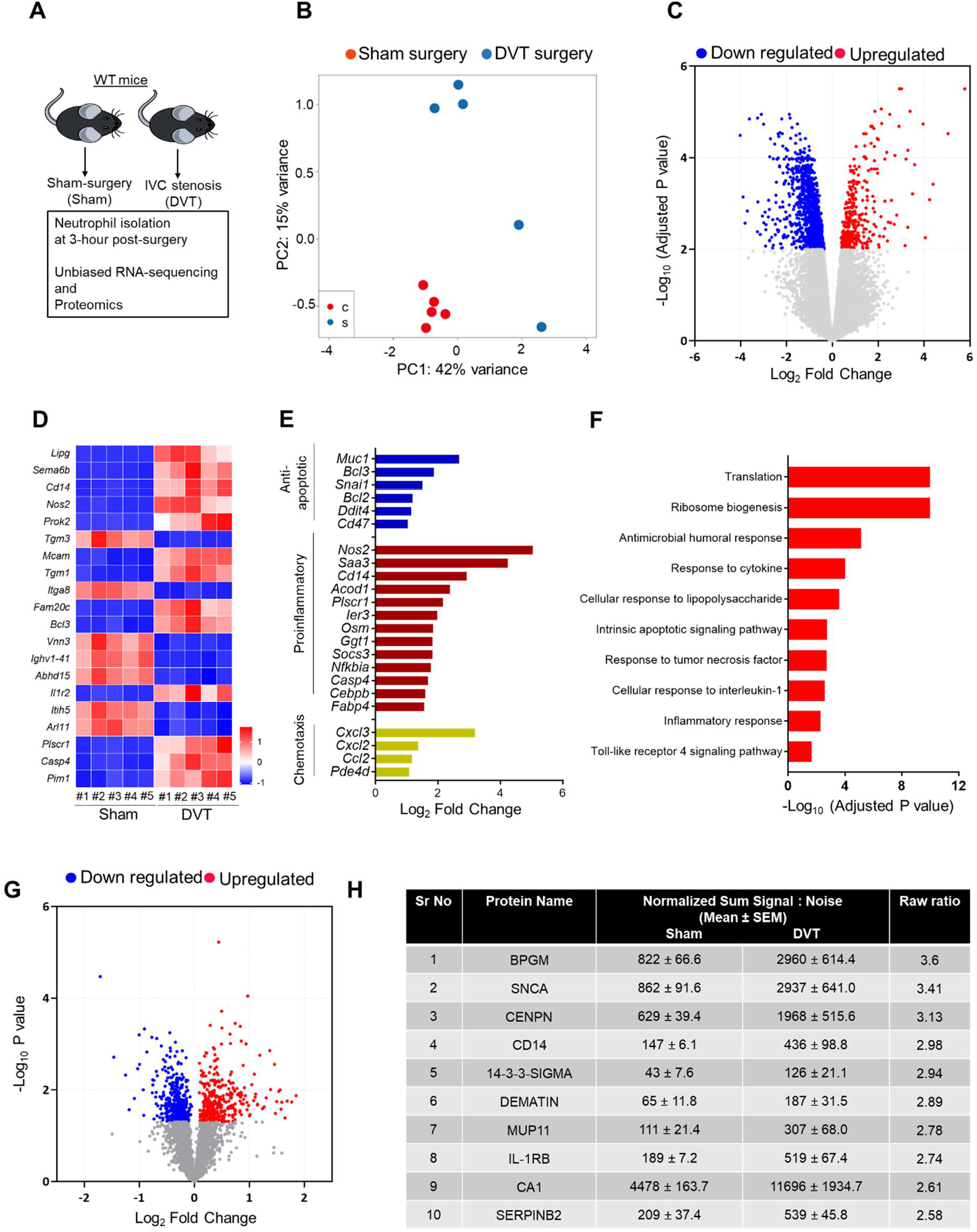
Transcriptomics and proteomics analysis of post-DVT neutrophils revealed aberrant overexpression of CD14. (A) Schematic of experimental design. (B) Principal component analysis was performed based on RNA-sequencing of neutrophils isolated from mice with sham-surgery or DVT. (C) Volcano plots of differentially expressed genes (DEGs) based on RNA-sequencing comparing neutrophils isolated from mice with sham-surgery or DVT. (D) Heatmap of top-20 DEGs. (E) Log fold-change of selected genes from DEGs (F) Pathways enriched in the upregulated DEGs based on gene ontology enrichment analysis. (G) Volcano plots of differentially expressed proteins based on proteomics data comparing neutrophils isolated from mice with sham-surgery or IVC stenosis. (H) top-10 upregulated genes in mice with IVC stenosis. n = 5 (B-F), n = 4 (G-H).

To assess whether the transcriptional alternations are consistent at the protein level, we applied proteomics on neutrophils isolated 3-hour post-surgery from mice with IVC stenosis and sham surgery. The analysis yielded 4645 proteins in total, among which 764 proteins were differentially regulated when comparing the IVC stenosis to the sham-surgery group (P<0.05, **Figure 2G**). The top 10 significantly upregulated and downregulated proteins are shown in **Figure 2H** and **Supplemental Fig 3**, respectively. In accordance with the RNA-seq analysis, neutrophils from mice with IVC stenosis exhibit increased levels of proteins associated with inflammation (CD14, SERPINB2, ALOX15, JDP2, NFkB-p100, SZRD1, and CREB).

### CD14 blockade did not improve DVT severity following IVC stenosis in mice

Considering that both the RNA-seq and proteomics analyses indicated CD14 is overexpressed in neutrophils from mice with IVC stenosis, we next focused on the role of CD14 in DVT outcomes. CD14 is a glycosylphosphatidylinositol-anchored membrane protein, a co-receptor for TLR4, that is known to be highly expressed in monocytes and at a lower level in neutrophils. Since CD14 is known to expressed on both, monocytes and neutrophils, to determine in which cell type CD14 is overexpressed post-IVC stenosis, we performed a flow cytometric analysis of CD14 in circulating monocytes and neutrophils, which showed that CD14 was significantly increased in neutrophils but not in monocytes (**Supplemental Figure 4**).

Next, we evaluated the therapeutic potential of targeting CD14 using a specific anti-CD14 antibody, 4C1. This antibody is a well-characterized CD14-blocking antibody that has been shown to have an inhibitory effect on endotoxin-induced acute lung injury and immune responses against gram-negative bacteria in mouse models.^25,26^ Male WT mice were randomly assigned to receive either 4C1 or isotype control antibody (IgG2b, κ) (4 mg/kg intravenously, 30 min before surgery and 24 h post-surgery), and DVT severity was evaluated 48 hours following IVC stenosis. CD14 blockade did not have a significant effect on DVT severity as evaluated by IVC thrombus weight, thrombus length, thrombosis incidence, or plasma elastase levels (P> 0.05 vs isotype control antibody treated mice, **Figure 3A-B**). These data are in contrast to a previous study suggesting that CD14 inhibition improves survival and attenuates thromboinflammation in a baboon model of *Escherichia coli* sepsis. This suggests that CD14 inhibition may be protective against thrombosis during sepsis-like conditions, but not during sterile inflammation, such as DVT. A previous report suggested that CD14 deficiency resulted in reduced production of IL-10,^27^ a potent anti-inflammatory cytokine. IL-10 is also known to modulate thrombus-induced vein wall inflammation and venous thrombosis.^28^ To investigate the effect of CD14 blockade, we evaluated plasma IL-10 levels in mice with IVC stenosis treated with 4C1 and isotype control antibody. We observed a significant reduction in plasma IL-10 levels in mice treated with the 4C1 antibody (P<0.05 vs isotype control antibody-treated mice, **Figure 3D)**. These data suggest that any positive effects of CD14 blockade on thrombus propagation may be counterbalanced by a significant decrease in IL-10 levels, resulting in no net change in DVT severity.

**Figure 3:**
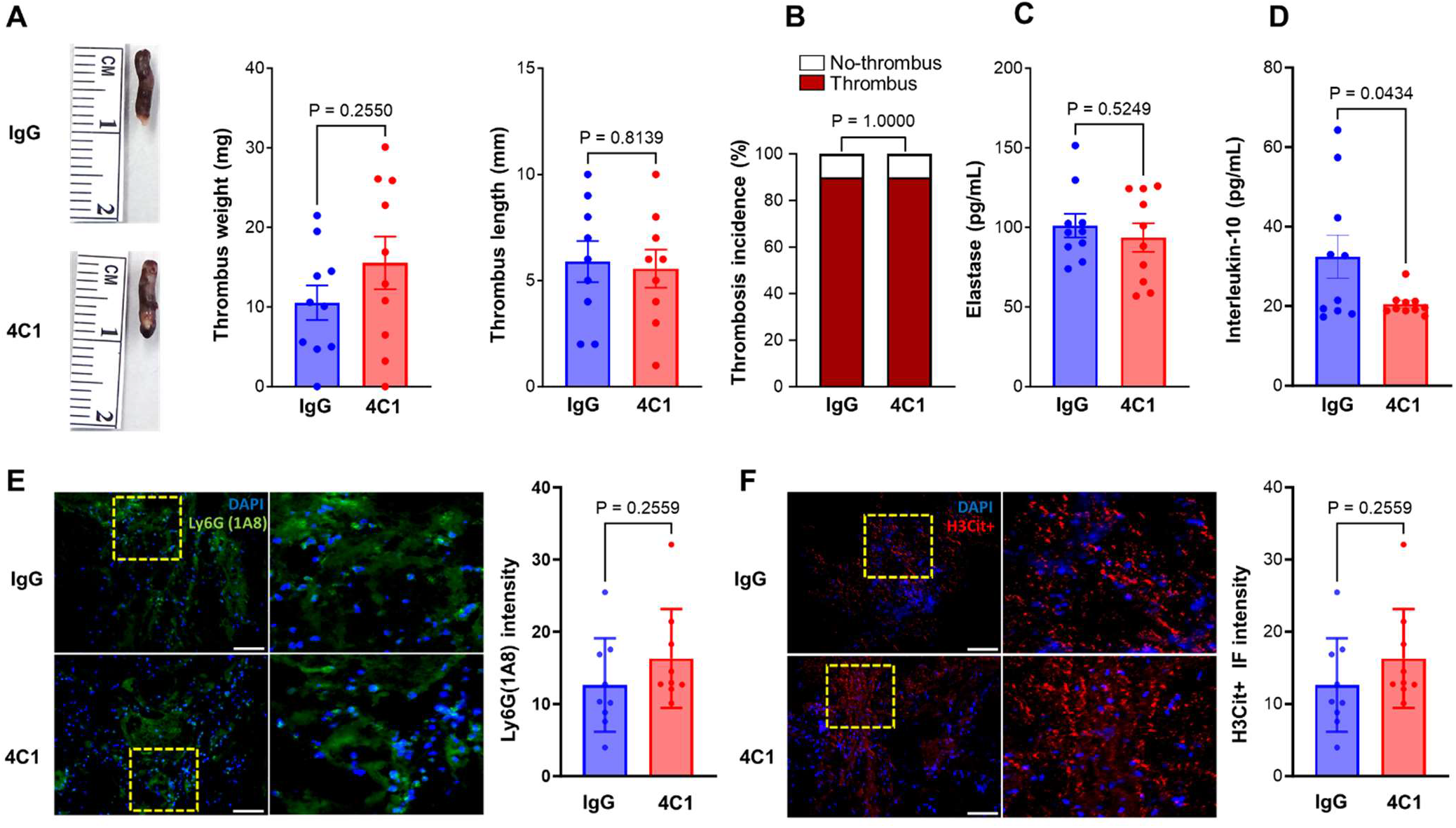
CD14 blockade did not improve DVT severity following IVC stenosis in mice. (A) Left, representative IVC thrombus harvested 48-hour post-stenosis from each group. Middle, thrombus weight (mg), right, thrombus length (mm). (B) Thrombosis incidence. Plasma samples were prepared 48-hour post-stenosis from each group. (C) Plasma levels of elastase and (D) plasma levels of IL-10 from each group. (E) Left, representative cross-sectional immunofluorescence image of the isolated IVC (on day-2 post-stenosis) from each group for Ly6G (neutrophils, green) and (F) anti-histone H3 (citrulline R2 + R8 + R17) (NETs, red) and DAPI (blue). Magnification 20X; Scale bar 100 μm. Right, Quantification of the % Ly6G and % CitH3 positive cells. Data are mean ± SEM and analyzed by a Mann Whitney test (A) and an unpaired t test (C-F); Fisher’s exact test (B) n = 10 (A-F).

Recent studies have suggested that neutrophils are recruited to the IVC thrombus and contribute to the pathophysiology of DVT. To determine whether CD14 promotes neutrophil influx in the IVC thrombus following stenosis, we performed immunohistochemical analysis of the IVC thrombus. CD14 blockade with 4C1 did not result in significant changes in neutrophils or citrullinated histone H3-positive cells (a marker of NETs) within the thrombi (P<0.05 vs isotype control antibody-treated mice, **Figure 2E-F**). These data are in agreement with a previous report suggesting that neutrophil recruitment within the liver was completely independent of CD14 in LPS-treated mice.^29^

Our data provides unique insights into the transcriptional and proteomic changes in neutrophils during early IVC stenosis, which may contribute to venous thrombus propagation and DVT severity. These unbiased analyses shed new light on our understanding of neutrophil biology following DVT, and that may facilitate the development of effective therapeutics for DVT patients. Despite its strengths, our study has limitations. For example, in human blood, neutrophils represent the majority of white blood cells but are less abundant in mouse blood.

Moreover, certain cytokines and chemokines are differentially expressed in mice and humans. Thus, while we did not find any benefits of CD14 blockade on DVT outcomes in mice, it is possible that inhibition of CD14 in humans may exert positive effects on DVT outcomes. Nevertheless, we speculate that such a scenario is unlikely because of a recent phase 2 trial demonstrating that CD14 blockade did not improve the time-to-resolution of illness in hypoxemic patients with COVID-19.^30^

In summary, the current study supports the notion that IVC stenosis results in increased plasma G-CSF levels, neutrophilia, and neutrophil hyperactivation. Moreover, neutrophils isolated post-IVC stenosis exhibit overexpression of CD14. Nevertheless, blockade of CD14 with the monoclonal antibody 4C1 does not improve DVT severity in mice following IVC stenosis, suggesting a limited role of CD14 inhibition in reducing DVT severity.

## 4. RELATIONSHIP DISCLOSURES

The authors report nothing to disclose.

## ACKNOWLEDGMENTS

The authors acknowledge support from the National Institutes of Health (HL158546 to ND, HL150233, DK134011, and DK136685 to OR, HL098435, HL133497, HL141155 to AWO), Career Development Award from American Heart Association (20CDA3560123 to ND), and the American Heart Association postdoctoral fellowship (HK). We would like to acknowledge the services offered by the Center of Applied Immunology and Pathological Processes Immunophenotyping Core supported by the NIH CoBRE Award P20 GM134974 (PI, Andrew Yurochko). We would like to acknowledge the services offered by the Flow Cytometry Core of the Louisiana State University Health Sciences Center-Shreveport.

## AUTHOR CONTRIBUTIONS

NP, HK, LC, SKA, performed experiments, analyzed data, and co-wrote the manuscript; HC, TM, AWO, KYS and OR interpreted the data, and co-wrote the manuscript; and ND designed the research, analyzed, and interpreted the data, and wrote the manuscript.

## SUPPLLIMENTAL MATERIAL

### METHODS

#### RNA sequencing and data analysis

Neutrophil cell-pellets were lysed, and RNA was isolated using the RNeasy Mini Kit (Qiagen #74106) as per manufacturer’s instructions. Libraries were prepared with the Stranded mRNA Prep, Ligation Kit (Illumina). One µg of RNA was processed for each sample and mRNA was purified and fragmented. cDNA was synthesized, and 3’ ends were adenylated. Anchor sequences were ligated to each sample and a limited-cycle PCR was performed to amplify and index the libraries. The average library size was determined using an Agilent TapeStation D1000 assay (Agilent Technologies) and libraries were quantitated with qPCR (Bio-rad CFX96 Touch Real-Time PCR, NEB Library Quant Kit for Illumina). Libraries were normalized to 0.5 nM and pooled. The library pool was denatured and diluted to approximately 100pM. A 1% library of 2.5pM PhiX was spiked in as an internal control. Paired end 76 x 76 base pair sequencing was performed on an Illumina NovaSeq 6000. Primary analysis, including base calling and quality scoring, was performed onboard the Illumina NovaSeq 6000 (NovaSeq Control Software v1.8.0; RTA v3). Samples were de-multiplexed, the adapter sequences were removed (the first 9 cycles of sequencing were trimmed), and FASTQ files were generated. Data analysis was performed as we previously described. The quality of the raw FASTQ files was checked through FastQC v0.11.8 (https://www.bioinformatics.babraham.ac.uk/projects/fastqc/). Trimmomatic v.0.35 was used to trim the low-quality reads with the parameters: SLIDINGWINDOW:4:20 MINLEN:25. The resulted high-quality reads were then mapped to the mouse reference genome (GRCm38.90) using STAR 2.5.1b11 and quantified by RSEM 1.2.3112 following the ENCODE-DCC/RNA-seq pipeline. Differentially expression analysis was performed using Bioconductor limma + Voom (Version: 3.48.3) and EdgeR (Version: 3.34.1) packages.13,14 Benjamini-Hochberg correction was used to obtain adjusted p-values. Statistically significant differentially expressed genes were filtered based on adjusted p< 0.01. Genes with adjusted P value < 0.01 were considered as significant DEGs. The upregulated and downregulated DEGs were analyzed for significantly enriched Gene Ontology pathways using the clusterProfiler package (version 4.8.3). The significance of the enrichment was determined by right-tailed Fisher’s exact test followed by Benjamini-Hochberg multiple testing adjustment. The heatmap of all shared DEG between stimulated vs unstimulated and KO vs control (Figure 4H) was generated by ComplexHeatmap R package (version 2.16.0). Hierarchical clustering of top 100 differentially expressed genes (supplementary figure 6 A and B) was visualized as a heatmap using R package gplots (version 3.1.3).

### Peptide and protein identification

Protein identification was evaluated by commercial service from Cell Signaling Technology (CST®), TMT10plex™ Total Proteome profiling service, that provides accurate global profiling of protein abundance in cells and uses multiplexed sample labeling with Tandem Mass Tags™ (TMT™) and liquid chromatography tandem mass spectrometry (LC-MS/MS). Mass spectra were evaluated by CST® using SEQUEST and the GFY-Core platform (Harvard University). Searches were performed against the 20180718 update of the Uniprot Homo sapiens database with a mass accuracy of ± 50 ppm for precursor ions and 0.02 Da for product ions. Total proteome data were filtered to a 1% peptide-level false discovery rate (FDR) with mass accuracy ± 5 ppm on precursor ions and presence ions; IMAC data were filtered to samples with a phosphorylated residue prior to filtering to a 1% protein-level FDR. All IMAC quantitative results were generated using Skyline 16 to extract the integrated peak area of the corresponding peptide assignments. Accuracy of quantitative data was ensured by manual review in Skyline or in the ion chromatogram files. TMT quantitative results were generated using the GFY-Core platform (Harvard University).

### Immunofluorescence staining

IVC thrombi were harvested 48-hour post-stenosis, detached from the vessel wall, dried and weighed in a microbalance followed by embedded in optimal cutting temperature compound, frozen at −80°C, and was cut with a cryotome (CryoStar NX70 Cryostat; ThermoFisher Scientific) into 10-μm sections. 10 µm cryosections were then blocked in a ready-to-use protein block (#ab64226, Abcam) for 30 min at room temperature followed by three times washing with 1X PBS-Tween (PBST). Sections were incubated overnight with primary antibodies for anti-rabbit Cit-Histone H3 (Arg2, Arg8, Arg17) (Thermo Fisher Scientific, #630-180ABBOMAX, 1:300) and FITC-Ly6G (Thermo Fisher Scientific, 11-9668-82, 1:250) at 4°C overnight. Next day, for anti-rabbit Cit-Histone H3 staining only, sections were washed with 1X PBST thrice and incubated with Alexa Fluor 647 goat anti-mouse (Thermo Fisher Scientific, #A21244, 1:500) secondary antibody for 1 h in the dark at room temperature. Slides were then washed three times with 1X PBST. All slides were then blotted dry, sealed with mounting medium containing DAPI (Thermo Fisher Scientific, #P36983) and allowed to dry overnight before imaging. Images were acquired using a fluorescence microscope EVOS™ M5000 Imaging System (Invitrogen) at 20 x magnification. Quantification was carried out using ImageJ software (NIH Image J, USA) and assessed with GraphPad Prism 8.0.0 to identify statistically significant differences among groups.

### Antibody treatment

Male WT mice were randomly assigned to receive either 4C1 (BD Pharmingen, purified NA/LE Rat Anti-Mouse CD14 # 557896) or isotype control antibody (BD Pharmingen Purified Rat IgG2b, κ Isotype Control # 553986) at the dose of 4 mg/kg intravenously, 30 mins before the surgery and 24 hours post-surgery.

### ELISA assay

Plasma G-CSF (R&D systems # MELA20), plasma IL-10 (R&D systems # M1000B-1), plasma elastase (R&D systems # MELA20) and plasma MPO (R&D systems # DY3667) were analyzed using commercially available kits.

## FIGURE LEGENDS

**Supplemental Figure 1:**
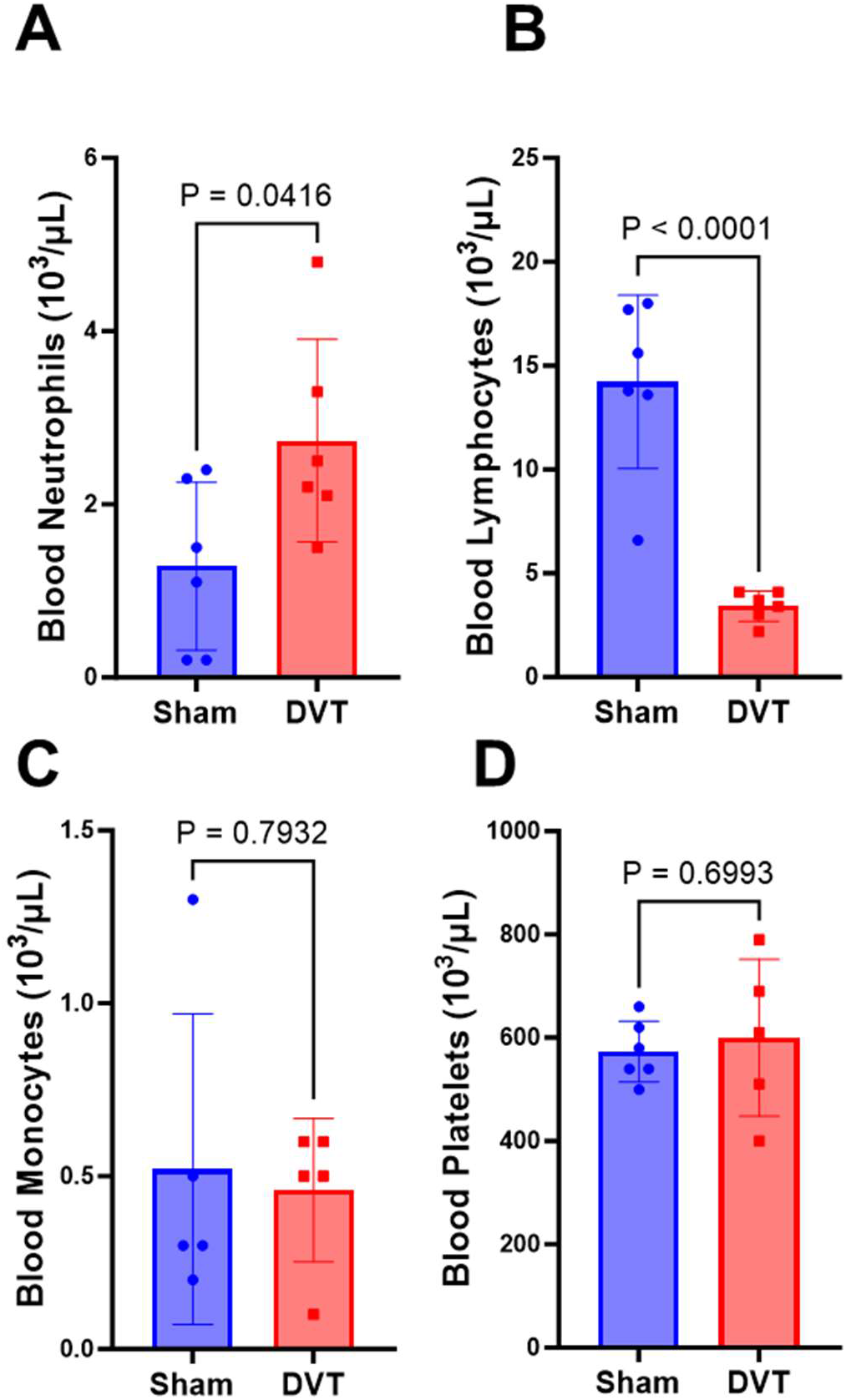
Blood was collected at 3-hr post-surgery from mice with sham-surgery and mice with DVT for analysis of blood neutrophil (A), blood lymphocytes (B), blood monocytes (C), platelets (D). All data are from male mice and are mean ± SEM and by an unpaired Student t test; n = 5 (A-D).

**Supplemental Figure 2:**
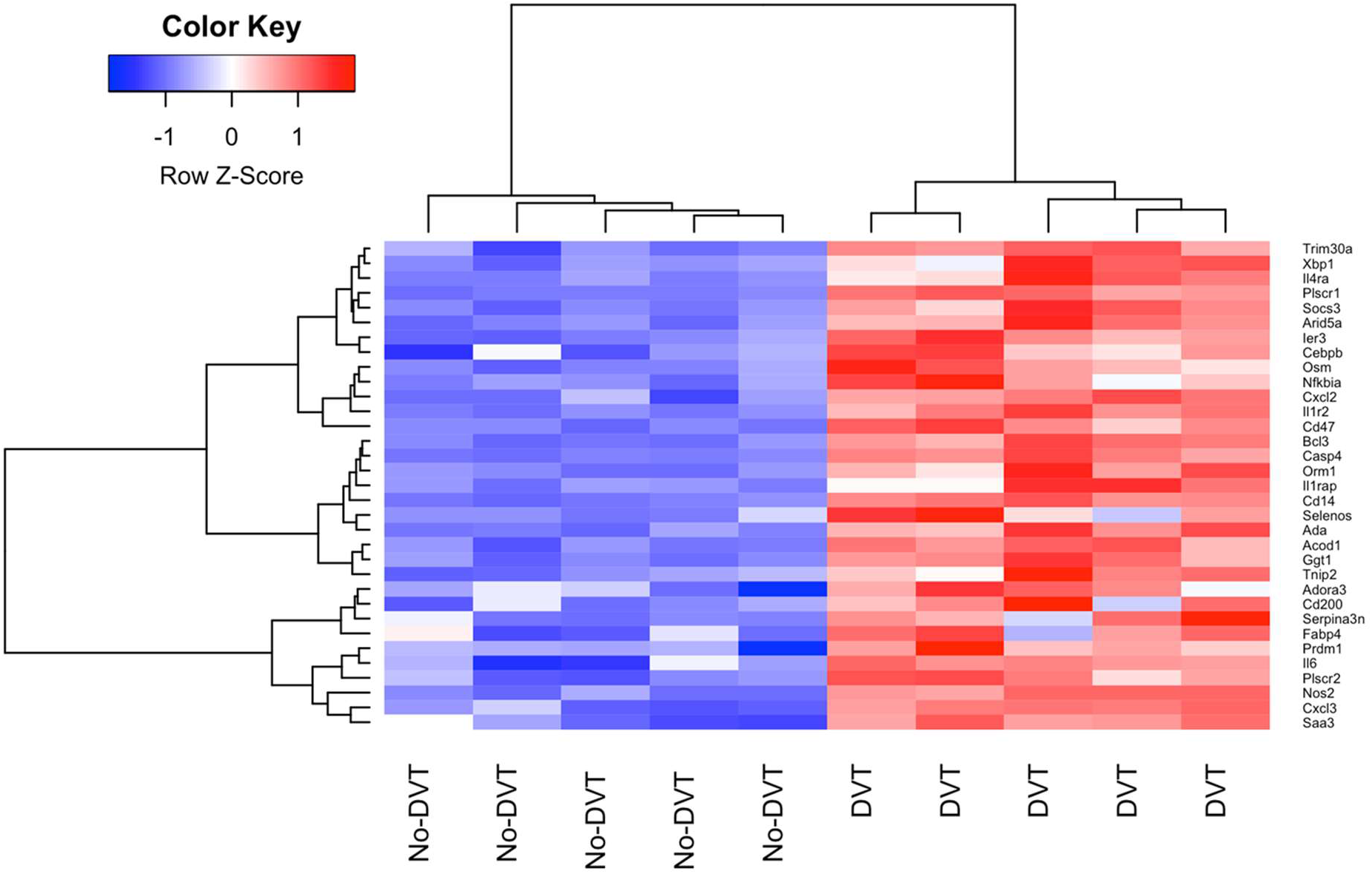
Hierarchical clustering of gene expression from RNA-seq data (3-hour post-surgery) of neutrophils isolated from mice with DVT and mice with sham-surgery.

**Supplemental Figure 3:**
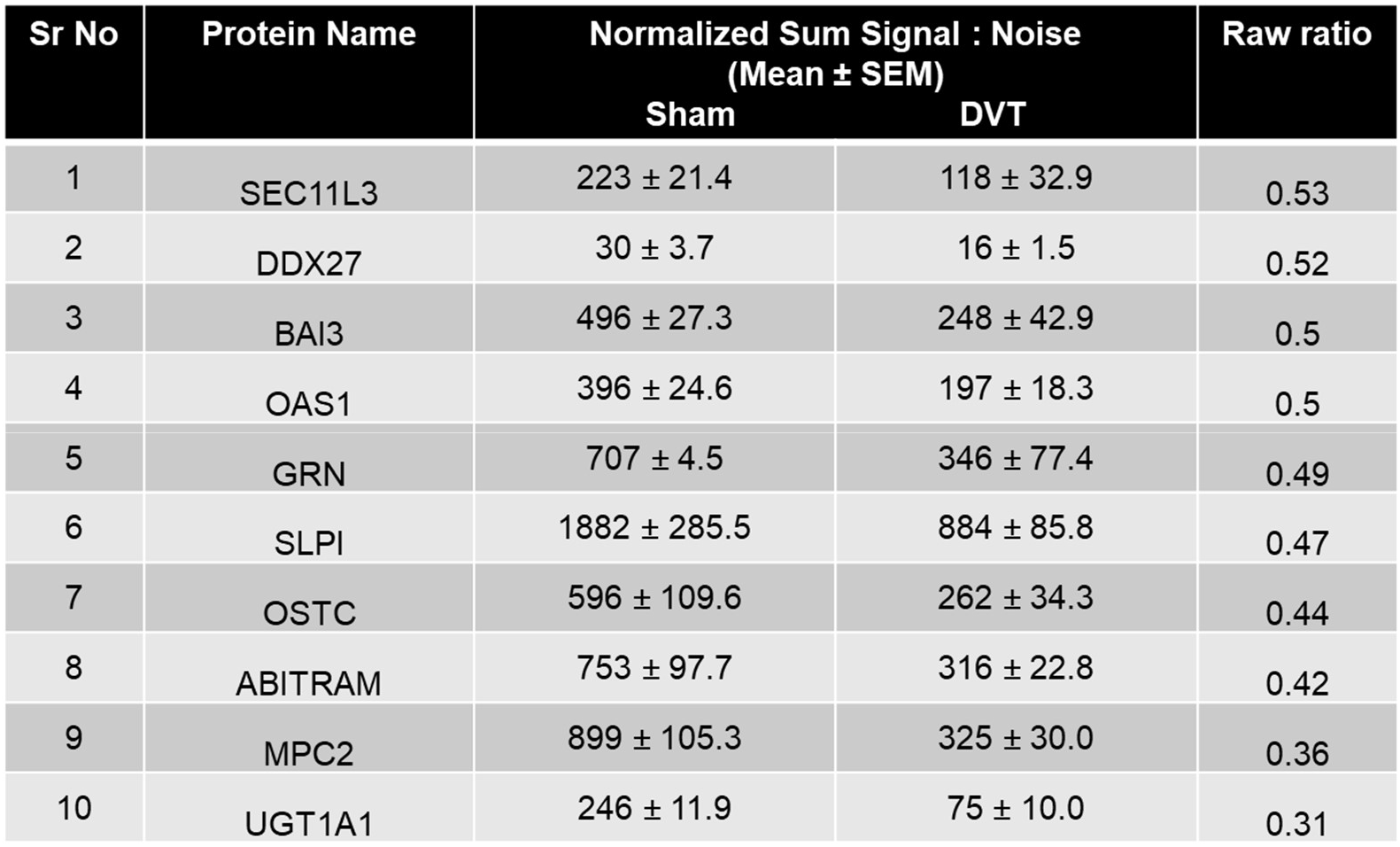
List of top 10 downregulated protein from neutrophils isolated from mice with DVT and mice with sham-surgery.

**Supplemental Figure 4:**
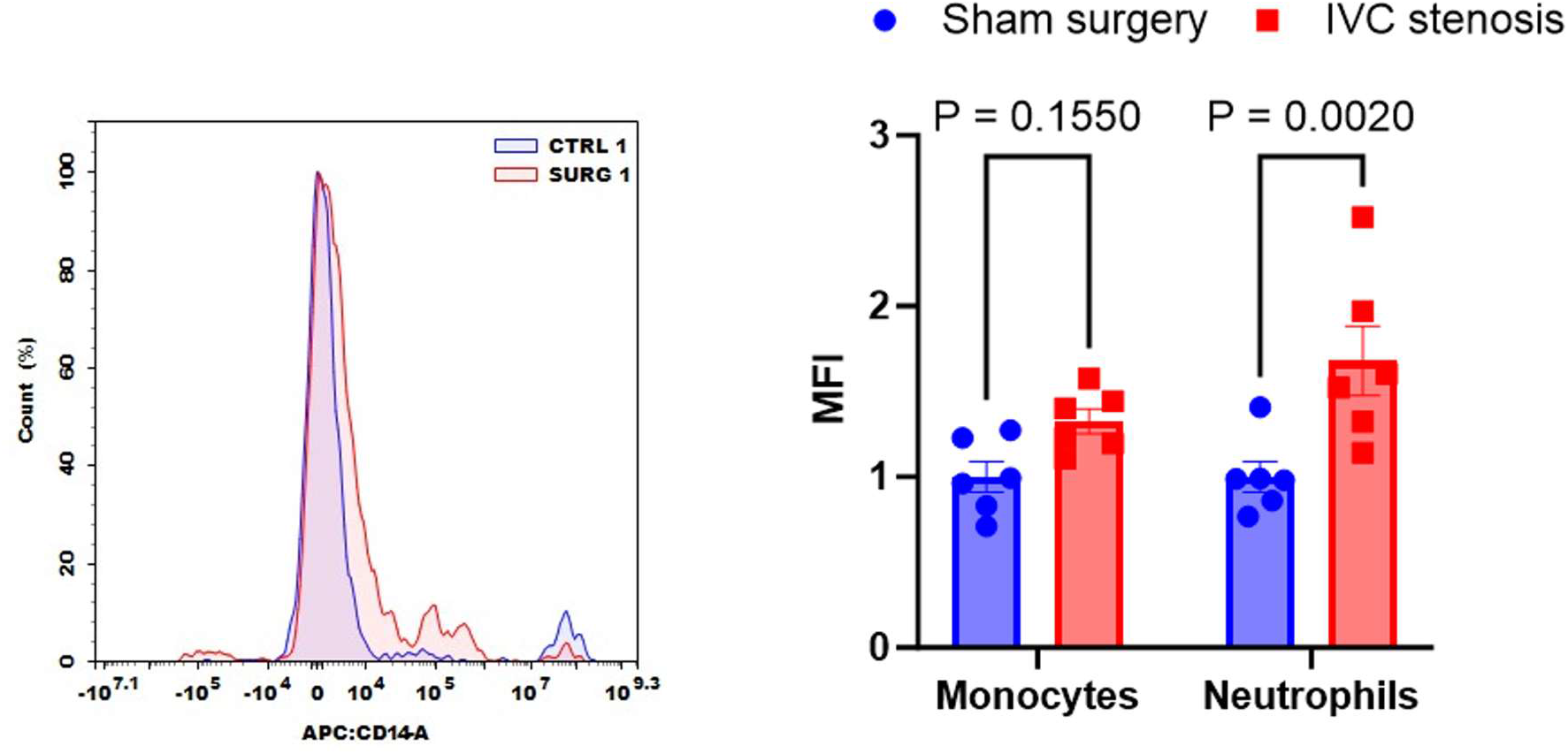
Left, representative image of flow-cytometric analysis of CD14 positive neutrophils. Right, quantification of CD14 expression in peripheral monocytes and neutrophils following IVC stenosis or sham-surgery in mice. Data are mean ± SEM, analyzed by two-way ANOVA followed by Holm-Šídák’s multiple comparisons test. n = 6.

## Notes

### Competing Interest Statement

The authors have declared no competing interest.

